# Somatic function of the Argonaute protein Aubergine is essential for neuromuscular development and function in *Drosophila*

**DOI:** 10.1101/2025.09.17.676627

**Authors:** P. Githure M’Angale, Devyn Oliver, Gimena Alegre, Jasmine Graslie, Angelina Tran, Alia Ohira, Max Zinter, Anna Malinkevich, Travis Thomson

**Affiliations:** Department of Neurobiology, University of Massachusetts Chan Medical School, Worcester, MA, 01605, USA; BPGbio, 300 3rd Avenue, Waltham, MA 02451; Worcester Technical High School, 1 Officer Manny Familia Way, Worcester, MA 01605

**Keywords:** Aubergine, Argonaute, PIWI, Copia, Transposable elements, Neuromuscular junction, Drosophila

## Abstract

**Background:** The PIWI-interacting RNA (piRNA) pathway is the primary defense against the deleterious activity of transposable elements (TEs), a role classically assigned to the germline. We recently discovered that the retrotransposon *Copia* is a negative regulator of synaptogenesis at the *Drosophila* larval neuromuscular junction (LNMJ) [1]. Here, we investigated whether the piRNA pathway regulates *Copia* in this somatic context.

**Methods:** Analysis of existing sequencing data revealed the expression of piRNA pathway components in somatic tissues [2]. We focused on Aubergine (*aub*), a core PIWI-clade Argonaute. We utilized CRISPR generated aub reporter lines and confocal microscopy to confirm the enrichment of AUB at the LNMJ and next generation sequencing coupled with digital PCR to validate the upregulation of TEs in *aub* knockdown larvae and adult tissues.

**Results:** Data from genetic reporters and antibody staining show that AUB is expressed and localized to the LNMJ. Tissue-specific knockdown of *aub* at the LNMJ resulted in increased TE expression, including *Copia*. In contrast to the synaptic overgrowth seen with *Copia* depletion [1], *aub* reduction caused a decrease in synapse number and impaired motor function and lifespan. These phenotypes are consistent with the upregulation of *Copia*, a negative regulator of synapse growth.

**Conclusions:** Our findings demonstrate that AUB functions somatically at the LNMJ to repress TEs, thereby ensuring proper neuromuscular development and function. This work establishes a physiological role for the piRNA pathway in a somatic tissue, linking TE repression to neuromuscular development.

## Background

Large swaths of eukaryotic genomes appear to have no function, in that they do not encode for canonical genes or structural elements needed for chromatin formation, transcription, and/or DNA replication. A significant portion of these genomic regions are comprised of transposable elements (TEs), which are obligate genome parasites. There is evidence that these regions do have a function as there can be strong selective pressure for their conservation; however, regarding TEs, it is often believed that such conservation is due to selective pressure on TEs to be effective parasites [3]. Recent research suggests that certain TEs have evolved functions in their host beyond their originally understood roles as parasites. Our group found that the *Drosophila* homolog of Arc (dArc1) is characteristic of a TE fragment and maintains many of its viral properties, including transferring across the synapse, and that this viral-like transfer is needed for structural synaptic plasticity [4]. Further we have found the retrotransposon Copia to be a potent regulator of structural synaptic plasticity in opposition to dArc1 [1].

Increased expression and transposition of TEs has been observed in mammalian neurons. For example, LINE-1 retrotransposon expression in mouse brains appears to be developmentally regulated and dynamic [5, 6]. However, the functional role of neuronally expressed TEs has remained elusive, and increased TE expression is observed in many neurological disorders. Modeling of Amyotrophic Lateral Sclerosis, Alzheimer’s disease, and Huntington’s disease in *Drosophila* suggests that the activation of TEs is the cause of some aspects of neurodegeneration [7]. A key phenotype consistent in these models is the presence of double-stranded DNA breaks, which previous works show as likely to be due to increased TE activity [7, 8].

Recently, we found that a spliced form of Copia is enriched in *Drosophila* motor neurons, and that Copia is a negative regulator of synaptic development, structural synaptic plasticity, and function [1]. While additional questions remain about how Copia regulates plasticity, one critical line of query is how Copia itself may be regulated to mitigate the genome damaging effects associated with transposition of TEs. Currently, the best understood mechanisms for the regulation of TEs is through small RNAs, specifically the PIWI-interacting RNA (piRNA) pathway. Most examples of the piRNA pathway are believed to be restricted to germline cells, or at most to somatic cells associated with the germline. However, there is growing evidence of piRNAs in somatic tissue and presence of somatic piRNAs in disease [7, 9–12]. Despite the apparent lack of piRNA pathway function in somatic cells, an interesting phenomenon has been observed upon mutation of the small interfering RNA (siRNA) pathway in *Drosophila* heads, in that a reduction of endogenous siRNAs lead not only to an increase in TE expression but also to an increase in what appear to be somatic piRNAs [13].

Key to the piRNA pathway are PIWI-type Argonautes Piwi, Aubergine (AUB), and Argonaute3 (AGO3). The phenotypes of PIWI-type Argonautes are mostly restricted to germline defects; however, there is a small but growing aggregate of evidence that mutations of piRNA pathway genes may influence neuronal development [14]. Also, there are many examples where a piRNA pathway gene can mitigate or exacerbate neurodegeneration phenotypes for diseases modeled in flies [14–16]. Specifically, the work reported in Bozetti et al. 2015 demonstrated that *aub* mutants have shorter LNMJ length, suggesting *aub* has a function in LNMJ development [17]. Thus, we opted to focus on *aub* directly at the *Drosophila* LNMJ, to determine whether Copia is a target of the piRNA pathway in somatic tissues.

In this paper we find that the PIWI-type Argonaute protein Aubergine (AUB) is needed for proper NMJ development in *Drosophila* larvae. We find that a specific spliced isoform of the retrotransposon Copia expression is increased both as an mRNA and protein at the *Drosophila* LNMJ. We previously found that *Copia* knockdown leads to an increase in synaptic boutons [1]. Here, we find that *aub* mutant flies exhibit an increase in Copia and a decrease in synaptic boutons, a phenotype consistent with Copia acting as a negative regulator of synaptogenesis.

## Results

### The Aub promoter is active in motor neurons

Owing to the role of Aub in *Drosophila* LNMJ development [17], we first tested if Aub is present in cells associated with the *Drosophila* LNMJ. To begin, we leveraged a CRISPR-mediated knock-in of Gal4 (CRIMIC, <u>CRI</u>SPR-<u>M</u>ediated Integration <u>C</u>assette) into the first intron of *aub*, allowing for endogenous *aub* promoter-driven expression of Gal4 [18, 19]. This expression when paired to the *Drosophila* Gal4/UAS system [20], to express a *UAS-mCD8::GFP* or *UAS-mCD8::RFP,* reports tissue and cell-type expression of *aub* (Fig. 1A).

**Figure 1.**
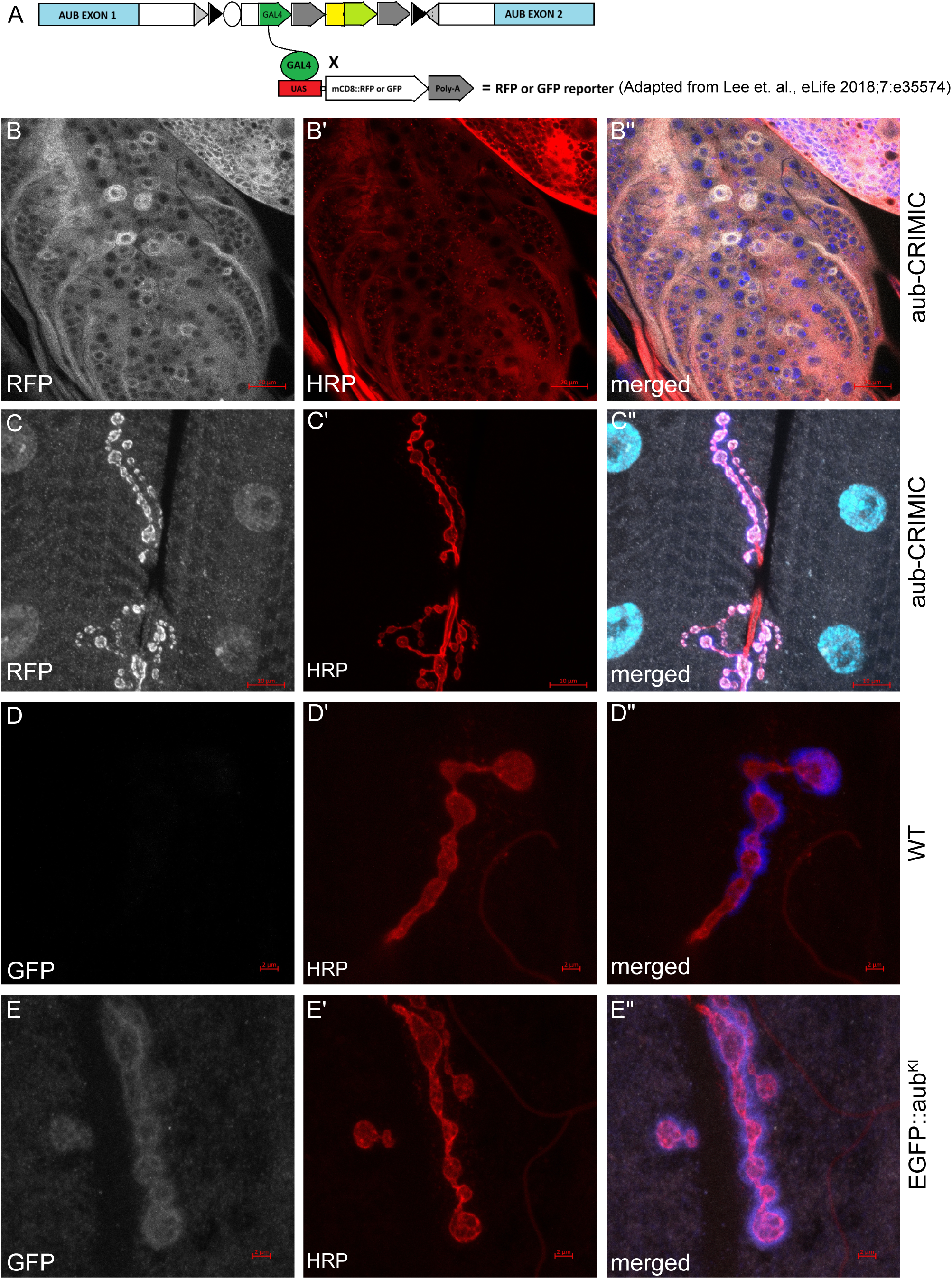
*Aub* expression is enriched in neuronal tissues. (A) Diagrammatic representation of the *aub*-CRIMIC fly line used to confirm *aub* expression. The endogenous *aub* promoter drives *Gal4* expression, which in turn leads to the expression of *RFP* or *GFP* under *UAS* control and results in RFP or GFP signal in all cells where *aub* is endogenously expressed. (B) Dissected third instar larval ventral nerve cord (VNC) of the *aub*-CRIMIC fly line, with RFP indicating the endogenous *aub* expression pattern in select motor neurons; (B’) horseradish peroxidase (HRP) staining of neuronal tissue; (B”) merged image of the orthogonal projection, showing both the endogenous *aub* expression pattern (RFP) and neuronal tissue staining (HRP). Scale bars, 20 μm. (C) Dissected third instar larval neuromuscular junction (LNMJ) of the *aub*-CRIMIC fly line, with RFP indicating the endogenous *aub* expression pattern at the LNMJ; (C’) HRP staining of the motor neurons of the LNMJ; (C”) merged image of the orthogonal projection, showing both endogenous *aub* expression pattern (RFP) and motor neuron staining (HRP). Scale bars, 10 μm. (D) Dissected third instar LNMJ of a WT (Canton-S) control fly line, with anti-GFP staining indicating lack of GFP-based signal in the control; (D’) HRP staining of motor neurons of the LNMJ; (D”) merged image of the orthogonal projection, showing negative GFP staining and positive motor neuron staining (HRP). Scale bars, 20 μm. (E) Dissected third instar LNMJ of the GFP knock-in *EGFP::aub^KI^* fly line, with anti-GFP staining indicating enrichment of full-length, GFP-tagged AUB at the LNMJ; (E’) HRP staining of motor neurons of the LNMJ; (E”) merged image of the orthogonal projection, showing both full-length AUB expression (GFP) and motor neuron staining (HRP). Scale bars, 2 μm.

Using this *aub-Gal4* reporter line we find that RFP is enriched in the ovary, in a pattern very similar to previously reported expression patterns of AUB in ovaries (Fig. S1A) [21]. There is also a cell-type specific pattern in larval brain lobes (Fig. S1B), the ventral nerve cord (Fig. 1B and Fig. S1C-D) and the LNMJ (Fig. 1C).

### AUB protein is enriched at the LNMJ

Using an existing CRISPR-Cas9-mediated GFP knock-in line, *EGFP::aub^KI^*[22], we confirm that this system has expression in the adult ovary similar to previously reported expression patterns for AUB [21] (Fig. S1F). We further find endogenous expression of EGFP-tagged AUB at the LNMJ (Fig. 1D-E and Fig. S1E and G) and is present in larval brain lobes (Fig. S1H-I). To determine if *aub* RNA itself was present at the LNMJ, we performed single molecule RNA-FISH and found that *aub* probes were enriched in the larval body wall muscle nuclei (Fig. S2A). We then generated antibodies against AUB, in rats and rabbits; we modeled our antibody production on previous works observing AUB in *Drosophila* ovaries [23]. With these antibodies we see a staining pattern in the adult *Drosophila* ovary consistent with previous publications (Fig. S2D-E). These AUB antibodies again show patterns (Fig. 2A-D) consistent with the CRIMIC and the EGFP::aub^KI^ patterns with enrichment at the LNMJ (Fig. S2F).

**Figure 2.**
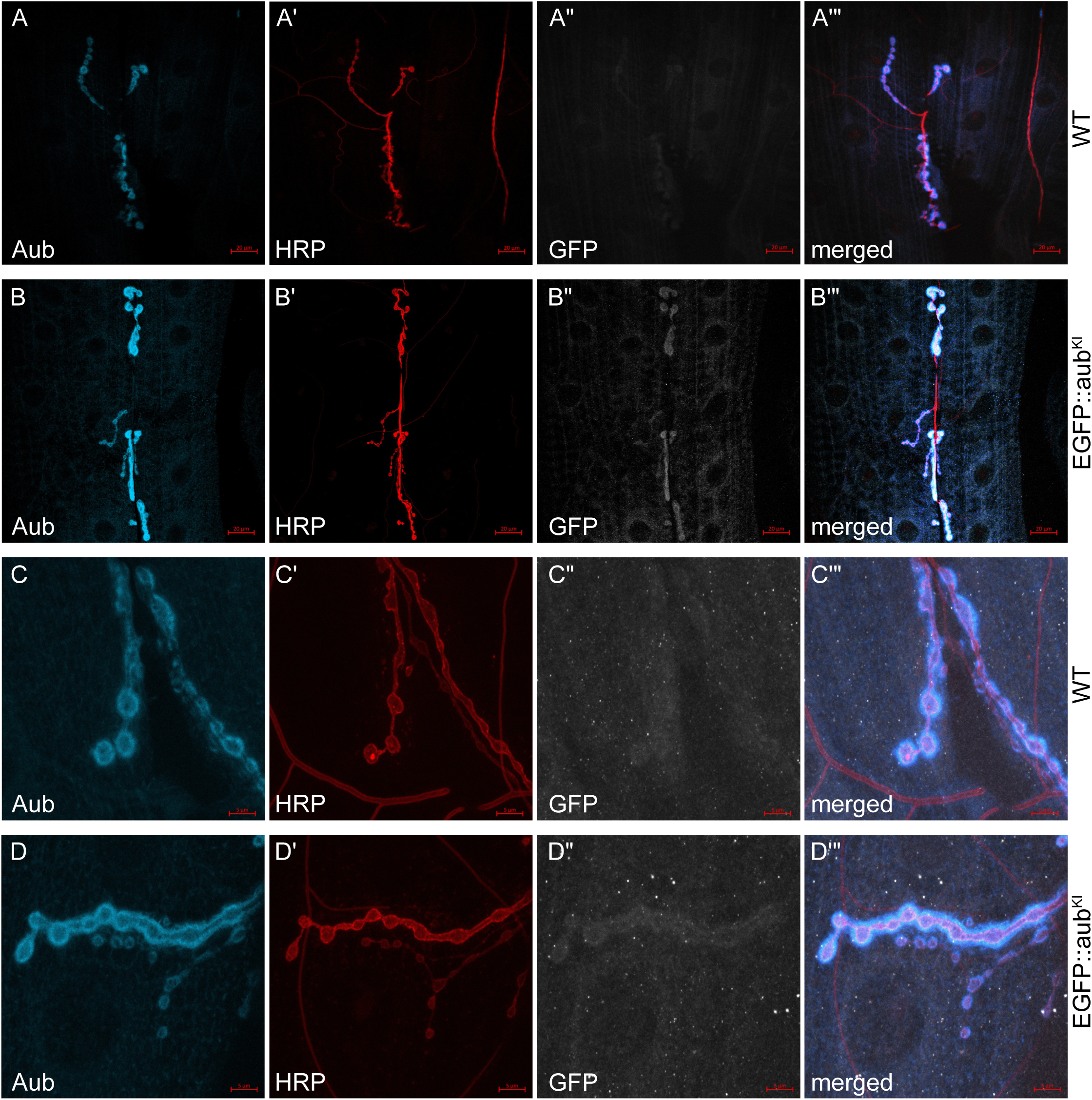
*Aub* protein is enriched in neurons and tissue around the LNMJ. (A) Dissected third instar larval body wall muscle (BWM) of a WT (Canton-S) control fly line, with anti-AUB staining indicating endogenous expression of full-length, WT *aub* protein; (A’) HRP staining of neuronal tissue; (A”) anti-GFP staining indicating lack of background GFP signal in the WT control; (A”’) merged image of the orthogonal projection. Scale bars, 20 μm. (B) Dissected third instar larval BWM of the GFP knock-in *EGFP::aub^KI^* fly line, with anti-AUB staining indicating expression of full-length, GFP-tagged AUB in a similar pattern of immunoreactivity as the WT *aub* protein expression observed in (A); (B’) HRP staining of neuronal tissue; (B”) anti-GFP staining indicating enrichment of full-length, GFP-tagged AUB in the same pattern of immunoreactivity as the anti-AUB staining pattern in (B); (B”’) merged image of the orthogonal projection. Scale bars, 20 μm. (C) Dissected third instar LNMJ of a WT (Canton-S) control fly line, with anti-AUB staining indicating endogenous expression of full-length, WT *aub* protein; (C’) HRP staining of motor neurons of the LNMJ; (C”) anti-GFP staining indicating lack of background GFP signal in the WT control; (C”’) merged image of the orthogonal projection. Scale bars, 5 μm. (D) Dissected third instar LNMJ of the GFP knock-in *EGFP::aub^KI^* fly line, with anti-AUB staining indicating expression of full-length, GFP-tagged AUB in a similar pattern of immunoreactivity as the WT *aub* protein expression observed in (C); (D’) HRP staining of motor neurons of the LNMJ; (D”) anti-GFP staining indicating enrichment of full-length, GFP-tagged AUB in a the same pattern of immunoreactivity as the anti-AUB staining pattern in (D); (D”’) merged image of the orthogonal projection. Scale bars, 5 μm.

### Reduction of Aub at the LNMJ shows a bilateral effect on Aub accumulation

We leveraged the *Drosophila* Gal4/UAS system to specifically reduce the expression of AUB in and around the LNMJ. Using either the motor neuron specific C380-Gal4 driver, or the muscle specific C57-Gal4 driver, we silenced *aub* by expressing two different long hairpins under UAS control generated by P-element insertion and phiC31 (UAS-aub-shRNAi^1^ and UAS-aub-shRNAi^2^ respectively) [20, 24]. Using this system, we can detect a decrease of *aub* mRNA by dPCR either pre-or postsynaptically (Fig. S2B-C). Expressing the aub-RNAi constructs in motor neurons, we see a reduction of AUB antibody staining at the LNMJ compared to wild type (WT) controls (Fig 3A-C). We quantified a substantial reduction of anti-AUB fluorescence in the presynaptic bouton (Fig 3A, D), while there is no reduction of AUB in the postsynaptic area (Fig 3A, E). Conversely, when *aub* is knocked down in the postsynaptic muscle, we see a reduction of AUB both in the pre-(Fig. 3H-J) and postsynaptic compartments (Fig. 3H-I, K).

**Figure 3.**
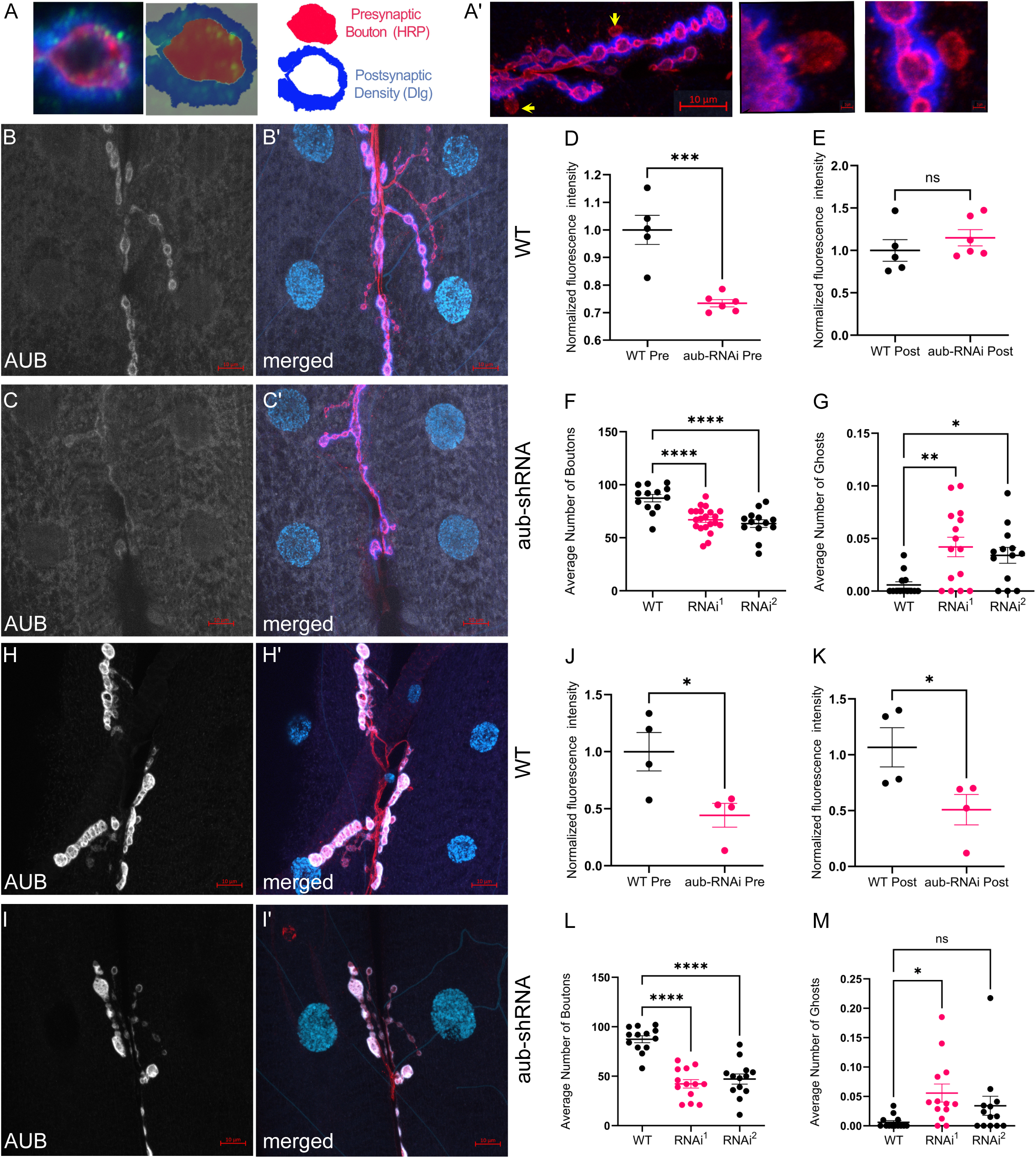
Neuronal *aub* knock down results in an LNMJ phenotype. (A) Diagrammatic representation and representative microscopy image examples of a standard bouton, highlighting the presynaptic bouton neuronal tissue (HRP staining) and postsynaptic bouton density (discs large (Dlg) antibody staining); (A’) representative microscopy image (scale bar 10 μm) examples of ghost boutons (yellow arrows, zoomed in boutons ((scale bar 1 μm))), which exhibit presynaptic neuronal tissue staining (HRP), but lack postsynaptic density staining. (B) Dissected third instar LNMJ of a WT (C380-Gal4/Canton-S) control fly line, with anti-AUB staining indicating endogenous expression of full-length, WT *aub* protein; (B’) merged image of the orthogonal projection with motor neuron staining (HRP). Scale bars, 10 μm. (C) Dissected third instar LNMJ of a motor neuron specific (C380-Gal4 driven) presynaptic *aub* knockdown fly line, with reduced anti-AUB staining indicating reduction of full-length, WT *aub* protein; (C’) merged image of the orthogonal projection with motor neuron staining (HRP). Scale bars, 10 μm. (D) Quantification of anti-AUB fluorescence intensity at the LNMJ, within the presynaptic bouton neuronal tissue; comparison of WT (C380-Gal4/Canton-S) control and motor neuron specific (C380-Gal4 driven) presynaptic *aub* knockdown fly lines. (E) Quantification of anti-AUB fluorescence intensity at the LNMJ, within the postsynaptic bouton density; comparison of WT (C380-Gal4/Canton-S) control and motor neuron specific (C380-Gal4 driven) presynaptic *aub* knockdown fly lines. (F) Quantification of standard bouton numbers in WT (C380-Gal4/Canton-S) control and two independent motor neuron specific (C380-Gal4 driven) presynaptic *aub* knockdown fly lines (RNAi^1^, RNAi^2^); N (left to right) = 13, 21 and 14. (G) Quantification of ghost bouton numbers in WT (C380-Gal4/Canton-S) control and two independent motor neuron specific (C380-Gal4 driven) presynaptic *aub* knockdown fly lines (RNAi^1^, RNAi^2^); N (left to right) = 15, 15 and 14. (H) Dissected third instar BWM of a WT (C57-Gal4/Canton-S) control fly line, with anti-AUB staining indicating endogenous expression of full-length, WT *aub* protein; (I’) merged image of the orthogonal projection with motor neuron staining (HRP). Scale bars, 10 μm. (I) Dissected third instar BWM of a muscle specific (C57-Gal4 driven) postsynaptic *aub* knockdown fly line, with reduced anti-AUB staining indicating reduction of full-length, WT *aub* protein; (I’) merged image of the orthogonal projection with motor neuron staining (HRP). Scale bars, 10 μm. (J) Quantification of anti-AUB fluorescence intensity in BWM, within the presynaptic bouton neuronal tissue; comparison of WT (C57-Gal4/Canton-S) control and muscle specific (C57-Gal4 driven) postsynaptic *aub* knockdown fly lines. (K) Quantification of anti-AUB fluorescence intensity in the BWM, within the postsynaptic bouton density; comparison of WT (C57-Gal4/Canton-S) control and muscle specific (C57-Gal4 driven) postsynaptic *aub* knockdown fly lines. (L) Quantification of standard bouton numbers in WT (C57-Gal4/Canton-S) control and two independent muscle specific (C57-Gal4 driven) postsynaptic *aub* knockdown fly lines (RNAi^1^, RNAi^2^); N = 13 for all genotypes. (M) Quantification of ghost bouton numbers in WT (C57-Gal4/Canton-S) control and two independent muscle specific (C57-Gal4 driven) postsynaptic *aub* knockdown fly lines (RNAi^1^, RNAi^2^); N = 13 for all genotypes. “ns” p≥0.05, * p<0.05, ** p<0.01, *** p<0.001, and **** p<0.0001. Error bars are SEM.

### Reduction of Aub at the LNMJ causes defects

The *Drosophila* LNMJ has been thoroughly analyzed for developmental defects and is a robust system that can be used to detect if a genetic perturbation is affecting aspects of synapse formation and/or function [25]. Synapses form in structures referred to as boutons, thus an increase or decrease in bouton number is a measure of synaptic development [26]. *Aub* mutant females have large increases in germline TE expression [27], and due to the viral-like nature of some TEs, these ovary derived TEs may travel throughout the larvae and affect somatic functions. To address this, we once again relied on our ability to perform cell-specific KD’s of *aub* in the presynaptic neuron (C380) (Fig. 3C) vs postsynaptic muscle (C57) (Fig. 3I) and observed a reduction of boutons (Fig. 3F and Fig. 3L, respectively). There is also an increase in a phenomenon referred to as “ghost” boutons, boutons that have the presynaptic label, HRP, but lack the postsynaptic marker, DLG (Fig. 3A’). The increased ghost bouton formation is observed with both presynaptic (Fig. 3G) and postsynaptic (Fig. 3M) knockdown. This suggests that a reduction of *aub* affects bouton formation as well as synapse maturation.

### Reduction of Aub in motor neurons results in decreased lifespan and motor functions

Owing to the somatic localization of Aub, and phenotypes at the LNMJ previously reported [17] and in this work, we tested if there are other phenotypes associated with a somatic reduction of Aub. *Aub* knockdown in either the neuron or muscle results in decreased crawling speed and crawling distance in third instar larvae (Fig. 4A-D).

**Figure 4.**
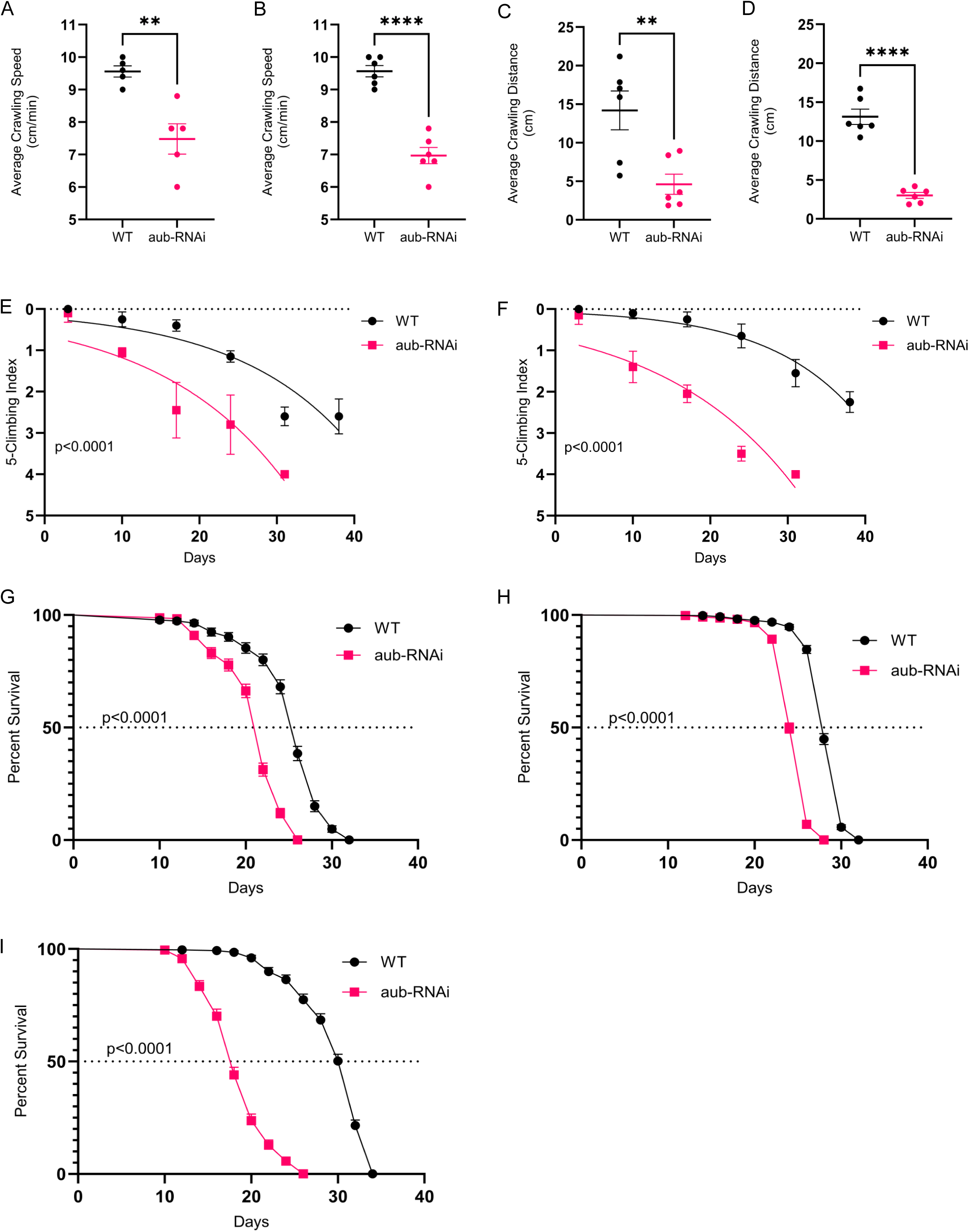
*Aub* knockdown in neurons and BWMs results in decreased lifespan and locomotor function. (A) Quantification of larval crawling speed, comparison of WT (C380-Gal4/Canton-S) control and motor neuron specific (C380-Gal4 driven) presynaptic *aub* knockdown fly lines; N = 50. (B) Quantification of larval crawling speed, comparison of WT (C57-Gal4/Canton-S) control and muscle specific (C57-Gal4 driven) postsynaptic *aub* knockdown fly lines; N = 60. (C) Quantification of larval crawling distance, comparison of WT (C380-Gal4/Canton-S) control and motor neuron specific (C380-Gal4 driven) presynaptic *aub* knockdown fly lines; N = 70. (D) Quantification of larval crawling distance, comparison of WT (C57-Gal4/Canton-S) control and muscle specific (C57-Gal4 driven) postsynaptic *aub* knockdown fly lines; N = 60. (E) Determination of locomotor ability, comparison of WT (C380-Gal4/Canton-S) control and motor neuron specific (C380-Gal4 driven) presynaptic *aub* knockdown fly lines; N = 50. (F) Determination of locomotor ability, comparison of WT (C57-Gal4/Canton-S) control and muscle specific (C57-Gal4 driven) postsynaptic *aub* knockdown fly lines; N = 50. (G) Analysis of lifespan, comparison of WT (C380-Gal4/Canton-S) control and motor neuron specific (C380-Gal4 driven) presynaptic *aub* knockdown fly lines; N = 266 (WT) and 243 (*aub* RNAi). (H) Analysis of lifespan, comparison of WT (C57-Gal4/Canton-S) control and muscle specific (C57-Gal4 driven) postsynaptic *aub* knockdown fly lines; N = 419 (WT) and 476 (*aub* RNAi). (I) Analysis of lifespan, comparison of WT (ELAV-Gal4/Canson-S) control and pan-neuron (Elav-Gal4 driven) presynaptic *aub* knockdown fly lines; N = 279 (WT) and 211 (*aub* RNAi). “ns” p≥0.05, * p<0.05, ** p<0.01, *** p<0.001, and **** p<0.0001. Error bars are SEM.

Due to the defects in LNMJ formation as well as larval crawling phenotypes, we wanted to address whether this has long-term physiological effects on the flies, specifically locomotor function and longevity. The drivers we use for knocking down Aub are effective in the adult central nervous system (CNS) and muscle [28–30]. Reducing *aub* expression throughout the life of adult fruit flies leads to a significant decrease in climbing ability and longevity, whether the reduction of Aub is in motor neurons (Fig. 4E, G) or muscle (Fig. 4F, H). Using a pan-neuronal driver (ELAV-Gal4) for knockdown of *aub* mRNA [28], we see a more substantial decrease in longevity than with the motor neuron specific driver (Fig. 4I).

### Aub repression leads to increased expression of transposons

Aub in the germline represses TEs, as such we wanted to measure TE expression in *aub* knockdown. RNA-sequencing data of brains isolated from larvae with *aub* KD in motor neurons reveals a change in expression of genes and a few specific seeds (genomic copies) of transposons (Fig. 5A). Mapping individual seeds of TEs is difficult due to the presence or absence of SNPs, insertions, or deletions and the redundancy of sequences of individual seeds. To address this, we also analyzed our data with a seed-agnostic approach, by mapping reads to consensus sequences of TEs (Fig 5B). We observe increases in expression of *Jockey*, *Copia*, *1731*, *TART-B*, and *Gypsy12*. In body wall muscle (BWM) where *aub* has been repressed using the C57-Gal4 driver we notice increases in specific seeds of *Copia* and *roo* (Fig 5C), and using the transposon-specific RNA-seq pipeline there is an observable increase in *Copia*, *invader1*, and *Juan* (Fig. 5D). We validated the increase in TE expression using dPCR, using probes and primers that are agnostic to the different seeds of TEs, and confirmed that Copia and other TEs are highly expressed in both pre-(Fig. 5E) and postsynaptic (Fig. 5F) knockdown of *aub*.

**Figure 5.**
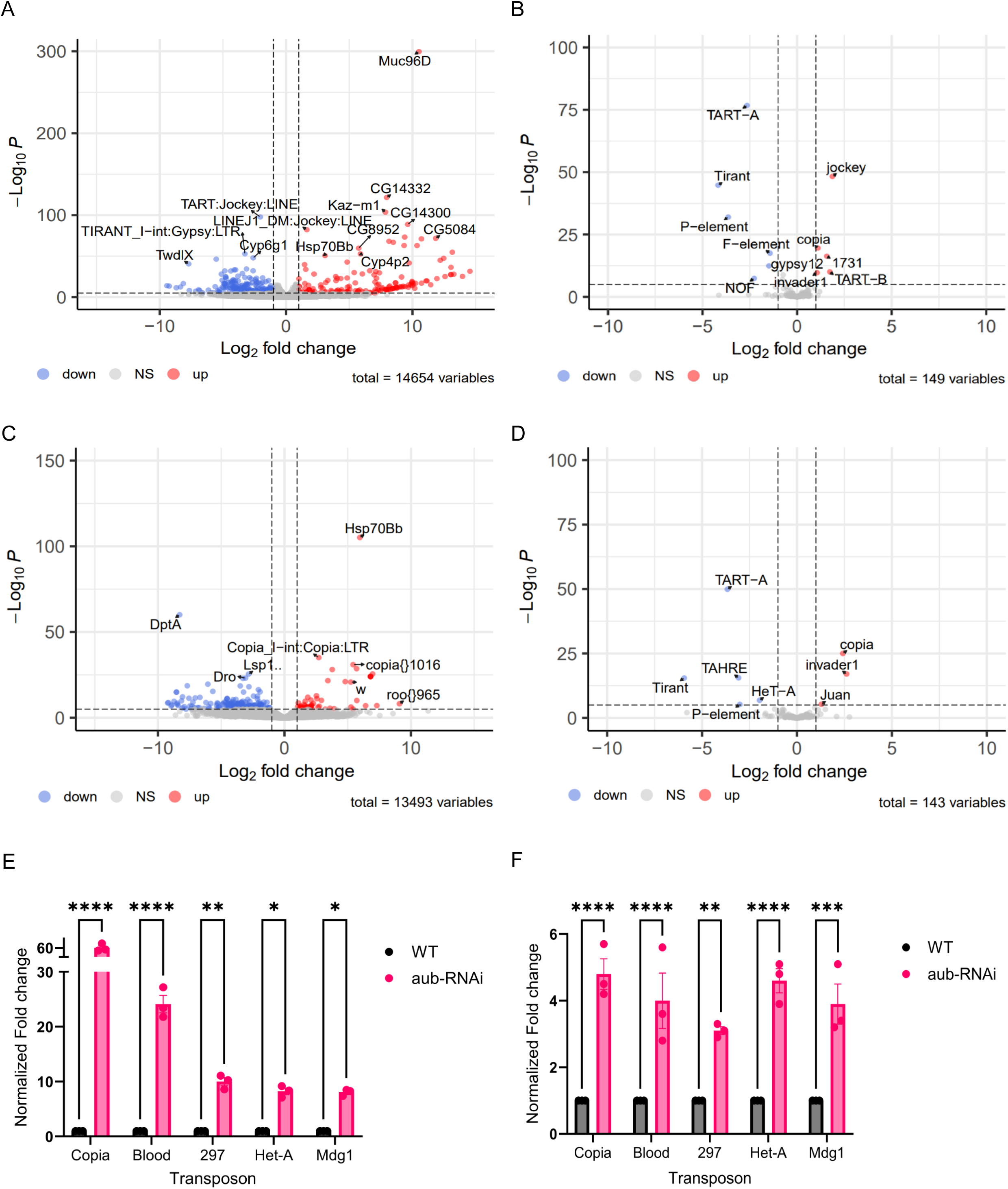
*Aub* knockdown in neurons and BWMs lead to differential expression of genes and transposons. (A) Volcano plot indicating differentially expressed transcripts in the larval central nervous system (CNS), as identified via RNA sequencing; comparison of WT (C380-Gal4/Canton-S) control and motor neuron specific (C380-Gal4 driven) presynaptic *aub* knockdown fly lines. Each dot represents one transcript; red dots denote significantly upregulated transcripts, and blue dots denote significantly downregulated transcripts, as detected in the *aub* knockdown fly line. (B) Volcano plot data as in (A), analyzed in a seed agnostic approach by using a consensus transposon-specific analysis pipeline. (C) Volcano plot indicating differentially expressed transcripts in the larval BWM, as identified via RNA sequencing; comparison of WT (C57-Gal4/Canton-S) control and muscle specific (C57-Gal4 driven) postsynaptic *aub* knockdown fly lines. Each dot represents one transcript; red dots denote significantly upregulated transcripts, and blue dots denote significantly downregulated transcripts, as detected in the *aub* knockdown fly line. (D) Volcano plot data as in (C), analyzed in a seed agnostic approach by using a consensus transposon-specific analysis pipeline. (E) Validation of increased expression of candidate TEs in the CNS, via dPCR; comparison of WT (C380-Gal4/Canton-S) control and motor neuron specific (C380-Gal4 driven) presynaptic *aub* knockdown fly lines. N = 3. (F) Validation of increased expression of candidate TEs in the BWM, via dPCR; comparison of WT (C57-Gal4/Canton-S) control and muscle specific (C57-Gal4 driven) postsynaptic *aub* knockdown fly lines. N = 3. Error bars are SEM.

### *Copia* mRNA and protein expression increases in *aub* mutants

As we observed an increase in *Copia* mRNA expression by RNA-seq and dPCR, we wanted to determine if the increase in *Copia* mRNA reflects an increase in Copia protein. Using two anti-Copia antibodies to stain larval body wall muscles, we tested if a knockdown of *aub* in the motor neurons or muscle leads to an increase of Copia protein. Anti-Copia^pol^, targets the polymerase region encoded by the full-length transcript (this paper), while anti-Copia^gag^, targets the Gag (capsid) protein [1]. In larvae with presynaptic motor neuron (C380-Gal4 driven RNAi) *aub* knockdown, immunostaining at the LNMJ with either antibody indicates an increase in Copia signal (Fig. 6A-C and Fig. 6D-F, respectively). Notably, there is also an increase in both Copia^gag^ and Copia^pol^ antibody staining at the LNMJ in larvae with *aub* knocked down in muscle (C57-Gal4 driven RNAi) (Fig. 6G-I and Fig. J-L, respectively).

**Figure 6.**
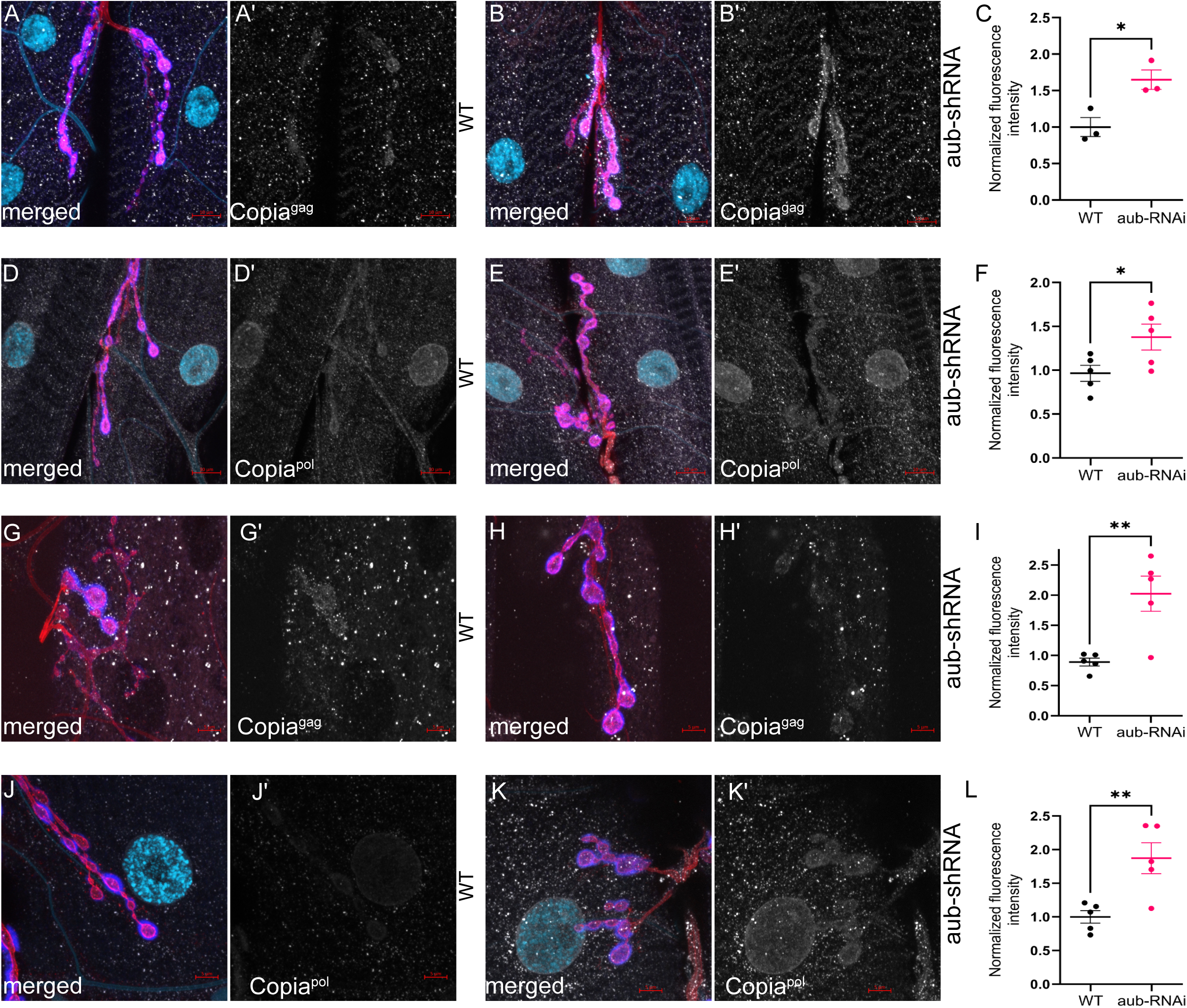
The retrotransposon Copia is enriched at the NMJs of *aub* knockdown larvae. (A) Orthogonal projection of dissected third instar LNMJ of a WT (C380-Gal4/Canton-S) control fly line, representing merged anti-Copia^gag^ staining and motor neuron staining (HRP); (A’) anti-Copia^gag^ antibody staining alone, indicating endogenous expression of the spliced isoform of the Copia protein (Copia^gag^). Scale bars, 10 μm. (B) Orthogonal projection of dissected third instar LNMJ of a motor neuron specific (C380-Gal4 driven) presynaptic *aub* knockdown fly line, representing merged anti-Copia^gag^ staining and motor neuron staining (HRP); (B’) anti-Copia^gag^ antibody staining alone, indicating expression of the spliced isoform of the Copia protein (Copia^gag^) in the *aub* knockdown line. Scale bars, 10 μm. (C) Quantification of Copia^gag^ protein levels via immunostaining analysis; comparison of WT (C380-Gal4/Canton-S) control and motor neuron specific (C380-Gal4 driven) presynaptic *aub* knockdown fly lines (A’ and B’, respectively). N = 3. (D) Orthogonal projection of dissected third instar LNMJ of a WT (C380-Gal4/Canton-S) control fly line, representing merged anti-Copia^pol^ staining and motor neuron staining (HRP); (D’) anti-Copia^pol^ antibody staining alone, indicating endogenous expression of the full-length isoform of the Copia protein (Copia^pol^). Scale bars, 10 μm. (E) Orthogonal projection of dissected third instar LNMJ of a motor neuron specific (C380-Gal4 driven) presynaptic *aub* knockdown fly line, representing merged anti-Copia^pol^ staining and motor neuron staining (HRP); (E’) anti-Copia^pol^ antibody staining alone, indicating expression of the full-length isoform of the Copia protein (Copia^pol^) in the *aub* knockdown line. Scale bars, 10 μm. (F) Quantification of Copia^pol^ protein levels via immunostaining analysis; comparison of WT (C380-Gal4/Canton-S) control and motor neuron specific (C380-Gal4 driven) presynaptic *aub* knockdown fly lines (D’ and E’, respectively). N = 5. (G) Orthogonal projection of dissected third instar LNMJ of a WT (C57-Gal4/Canton-S) control fly line, representing merged anti-Copia^gag^ staining and motor neuron staining (HRP); (G’) anti-Copia^gag^ antibody staining alone, indicating endogenous expression of the spliced isoform of the Copia protein (Copia^gag^). Scale bars, 5 μm. (H) Orthogonal projection of dissected third instar LNMJ of a muscle specific (C57-Gal4 driven) postsynaptic *aub* knockdown fly line, representing merged anti-Copia^gag^ staining and motor neuron staining (HRP); (H’) anti-Copia^gag^ antibody staining alone, indicating expression of the spliced isoform of the Copia protein (Copia^gag^) in the *aub* knockdown line. Scale bars, 5 μm. (I) Quantification of Copia^gag^ protein levels via immunostaining analysis, comparison of WT (C57-Gal4/Canton-S) control and muscle specific (C57-Gal4 driven) postsynaptic *aub* knockdown fly lines (G’ and H’, respectively). N = 5. (J) Orthogonal projection of dissected third instar LNMJ of a WT (C57-Gal4/Canton-S) control fly line, representing merged anti-Copia^pol^ staining and motor neuron staining (HRP); (J’) anti-Copia^pol^ antibody staining alone, indicating endogenous expression of the full-length isoform of the Copia protein (Copia^pol^). Scale bars, 5 μm. (K) Orthogonal projection of dissected third instar LNMJ of a muscle specific (C57-Gal4 driven) postsynaptic *aub* knockdown fly line, representing merged anti-Copia^pol^ staining and motor neuron staining (HRP); (K’) anti-Copia^pol^ antibody staining alone, indicating expression of the full-length isoform of the Copia protein (Copia^pol^) in the *aub* knockdown line. Scale bars, 5 μm. (L) Quantification of Copia^pol^ protein levels via immunostaining analysis; comparison of WT (C57-Gal4/Canton-S) control and muscle specific (C57-Gal4 driven) postsynaptic *aub* knockdown fly lines (J’ and K’, respectively). N = 5. “ns” p≥0.05, * p<0.05, ** p<0.01, *** p<0.001, and **** p<0.0001. Error bars are SEM.

### Knockdown of *aub* alters the splicing of transposons

The full-length *Copia* transcript (*Copia^full^*) has long been known to possess a splice junction [31], that results in the 2 Kb RNA product, *Copia^gag^*, which is largely comprised of the capsid encoding region with most of the Pol region spliced out (Fig. 7A). From sequencing data, we found there is an increase in *Copia^gag^*mRNA due to more splicing events in *aub* neuronal (Fig. 7B-C, F) and body wall muscle (Fig. 7D-E, G) knockdown lines. In wild type flies we find the ratio of *Copia^full^* vs. *Copia^gag^* is skewed towards *Copia^full^* [1]. However, in *aub* mutant LNMJs, *Copia^gag^* is present at a much higher ratio when compared to Copia^full^ (Fig. 7H).

**Figure 7.**
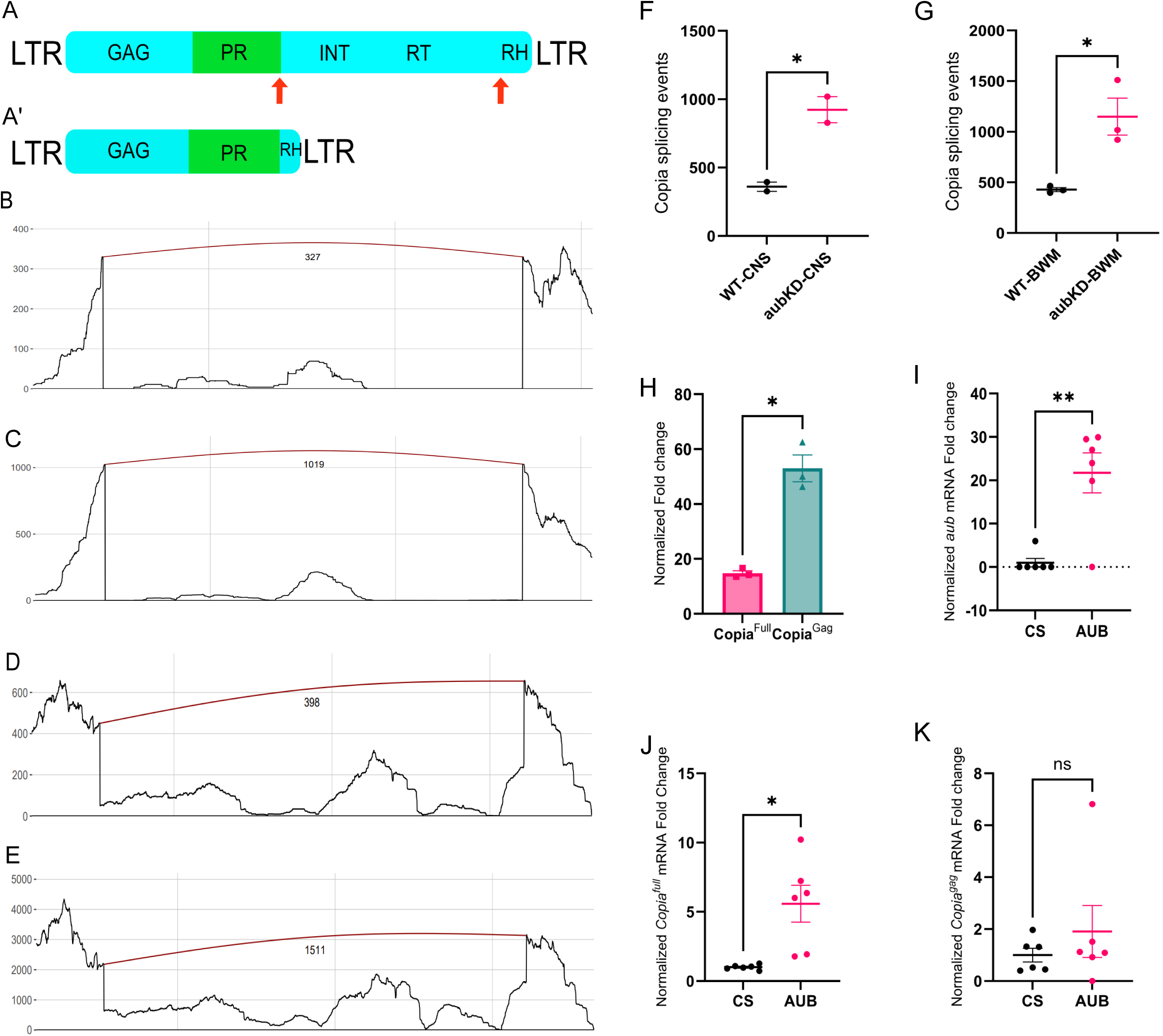
Misexpression of *aub* results in aberrant splicing of transposons. (A) Diagrammatic representation of the Copia retrotransposon full-length isoform (*Copia^full^*), with major structural regions noted; splicing at the red arrows generates the shorter *Copia^gag^* isoform. LTR, long terminal repeats; GAG, group antigens; PR, protease; INT, integrase; RT, reverse transcriptase; RH, RNase H. (A’) Diagrammatic representation of the Copia retrotransposon spliced isoform (*Copia^gag^*) product, resulting from splicing of the *Copia^full^* isoform in (A). (B) SpliceWiz coverage plot depicting alternative splicing events for Copia in CNS tissue, in a WT (C380-Gal4/Canton-S) control fly line. (C) SpliceWiz coverage plot depicting alternative splicing events for Copia in CNS tissue, in the motor neuron specific (C380-Gal4 driven) presynaptic *aub* knockdown fly line. (D) SpliceWiz coverage plot depicting alternative splicing events for Copia in BWM tissue, in a WT (C57-Gal4/Canton-S) control fly line. (E) SpliceWiz coverage plot depicting alternative splicing events for Copia in BWM tissue, in the muscle specific (C57-Gal4 driven) postsynaptic *aub* knockdown fly line. (F) Quantification of Copia splicing events in CNS tissue; comparison of WT (C380-Gal4/Canton-S) control and motor neuron specific (C380-Gal4 driven) presynaptic *aub* knockdown fly lines. N = 3. (G) Quantification of Copia splicing events in BWM tissue; comparison of WT (C57-Gal4/Canton-S) control and muscle specific (C57-Gal4 driven) postsynaptic *aub* knockdown fly lines. N = 3. (H) Quantification of alternative splicing of Copia in larval CNS tissue via dPCR; comparison of WT (C380-Gal4/Canton-S) control and motor neuron specific (C380-Gal4 driven) presynaptic *aub* knockdown fly lines. Copia^full^ probe recognizes unspliced, full-length Copia, as diagrammed in (A); Copia^gag^ probe recognizes spliced Copia, as diagrammed in (A’). N = 3. (I) Quantification of *aub* RNA IP from adult fly heads, utilizing the EGFP::aub^KI^ fly line; comparison to the WT (Canton-S) control. N = 6. (J) Quantification of *Copia^full^* RNA IP from adult fly heads, utilizing the EGFP::aub^KI^ fly line; comparison to the WT (Canton-S) control. N = 8. (K) Quantification of *Copia^gag^* RNA IP from adult fly heads, utilizing the EGFP::aub^KI^ fly line; comparison to the WT (Canton-S) control. N = 6. “ns” p≥0.05, * p<0.05, ** p<0.01, *** p<0.001, and **** p<0.0001. Error bars are SEM.

### AUB binds its own mRNA and the full-length *Copia* transcript

To determine if AUB’s regulatory effects are direct, we performed RNA immunoprecipitation (RIP) from adult fly heads using the *EGFP::aub*^KI^ line, followed by dPCR. We found that AUB binds to its own mRNA, a known feature of autoregulation in the germline [32] [33] (Fig 7I).

Importantly, AUB also specifically immunoprecipitated the full-length *Copia* transcript, but not the spliced *Copia^gag^* variant (Fig 7J-K). This indicates that AUB directly and selectively interacts with the unspliced, replication-competent form of *Copia* RNA.

## Discussion

Here we describe that the *Drosophila* PIWI-type argonaute, Aubergine, shows a somatic localization and functions to repress TEs at the *Drosophila* LNMJ. Specifically, we observed that (i) the *aub* promoter is active in motor neurons, (ii) *aub* mRNA is present in cells that comprise the LNMJ, (iii) AUB endogenously tagged with GFP localizes to the LNMJ, (iv) AUB antibodies also label the LNMJ and finally (v) the expression of *aub* shRNA constructs reduce AUB at the LNMJ. The reduction of AUB protein at the LNMJ affects synaptogenesis, consistent with previous data that *aub* is needed for proper LNMJ size [17]. Further, this reduction of Aub in motor neurons leads to an increase in TE expression in both motor neurons and muscles. Specifically, we find an increase in the retrotransposon Copia, at the LNMJ as measured by RNA-seq, dPCR, and Copia antibody staining. In addition to the decrease of synaptic boutons at the larval NMJ, when *aub* is knocked down pan-neuronally, we also observe slower larval crawling and decreased fly longevity. Altogether this is convincing evidence that a piRNA gene is expressed in somatic tissue, regulates transposons and has tractable phenotypes.

While the piRNA pathway has a well-described role in repressing transposons in the germline of many metazoan species, the utility and presence of piRNAs outside of the germline is less clear. Most evidence of somatic piRNAs is through the presence of piRNA like small RNAs, or piRNA mutants exacerbating or mitigating disease states [34–36]. This work shows strong evidence for the somatic localization and function of AUB. We find that upon presynaptic knock down of *aub* in the motoneurons no significant decrease in the postsynaptic pool of AUB protein. Interestingly, the postsynaptic knockdown of *aub* results in a reduction of the pre– and postsynaptic pool of AUB protein, implying that the presynaptic accumulation of AUB requires WT postsynaptic AUB expression. In addition, a reduction of AUB in this somatic tissue leads to an increase in transposon expression. This is consistent with the role of the piRNA pathway in repressing TEs in the germline and germline-associated somatic tissue.

Our attempt to determine if AUB was directly associated with piRNAs using endogenously GFP-tagged AUB did not recover any small RNAs other than ribosomal RNAs. An obvious next experiment is to attempt small RNA co-immunoprecipitations with anti-AUB antibodies, use another Tag than GFP, and/or scaling up future attempts. Through AUB immunoprecipitation, we determined that AUB binds to its own RNA and that of the full-length Copia transcript (Copia^full^) but not to that of the alternatively spliced Copia, Copia^gag^. This specific interaction between AUB and the RNA of Copia could be a mechanism of how AUB regulates Copia expression possibly through translation initiation [33]. Further work must be done to determine if other piRNA pathway genes are present at the LNMJ, notably *argonaute-3* and *piwi*, both of which have described roles in TE repression in association with AUB [37]. Additionally, it would be useful to determine which other small RNAs may be interacting with AUB in somatic tissues.

Of note, the decrease in synapse formation upon reduction of AUB is the opposite phenotype to that observed when Copia is decreased at the *Drosophila* LNMJ. We have found that Copia is a potent repressor of synaptogenesis, and the observed decrease in synaptogenesis when Copia levels are increased is consistent with our previous work [1]. Using dPCR on larval CNS and body wall muscles, we confirmed that the expression of other TEs was upregulated by several fold compared to control tissues. This phenomenon is not surprising as *aub* in the germline is well-known to be a potent regulator of TEs [23, 38–41]. The increase of Copia protein is not proportional to the increase in mRNA, which suggests that other TE-repression pathways are redundant to the piRNA pathway and/or to post-transcription pathways that actively regulate TEs. One key pathway is that of the endogenous siRNAs, which has been shown to repress TEs in *Drosophila* heads; when the endogenous siRNA pathway is suppressed there is an increase in piRNA-like small RNAs [13]. This suggests a redundancy between the two pathways (siRNA/piRNA).

## Conclusion

In summary, this study provides compelling evidence for a novel, somatic function of the core piRNA pathway gene, *Aubergine*, at the *Drosophila* neuromuscular junction. We demonstrate that AUB is essential for proper synapse development, motor function, and lifespan. The mechanism underlying these functions involves the direct repression of transposable elements, including the retrotransposon *Copia*, within a somatic context. Our findings significantly broaden the established role of the piRNA pathway’s machinery, showing that its function extends beyond germline defense to play a critical role in the development and maintenance of the nervous system. This work establishes a vital link between TE regulation by a piRNA pathway component and neuromuscular health, opening new avenues for understanding the somatic roles of this conserved pathway in development and disease.

## Methods and Materials

### Fly Stocks and Husbandry

Fly lines used in this study; P[GD11831]v30124 (VDRC_v30124) aka aub-RNAi^1^ [42], P[KK101252]VIE-260B (VDRC_v106999) aka aub-RNAi^2^ [42], aub^QC42^ cn1 bw1/CyO (BDSC_4968), aub^HN2^ cn1 bw1/CyO (BDSC_8517), Canton-S (1, BDSC), UAS-EGFP (BDSC_5431), UAS-GFP-aub (BDSC_42219), EGFP::aub^KI^ [22] (gift from Dr. Tatjana Trcek), C380-Gal4 [43], C57-Gal4 [43], and elav-Gal4 (BDSC_458).

Female third-instar larvae were used for all NMJ dissections. Derivative lines were generated using standard chromosomal balancers and leveraging recombination. Flies were raised on standard *Drosophila* fly media at 25°C. Crosses were maintained at either 25°C or 29°C when driven by Gal4 in incubators with 12h light and dark cycles.

### Generation of Antibodies

Aub antibodies were generated against an Aub antigen by immunizing rabbits and rats with the peptide MNLPPNPVIARGRGRG, a peptide used previously to make anti-Aub antibodies [23] (Genscript). Anti-Copia^Pol^ custom antibody was generated against an antigen derived from the Copia “POL” region with the sequence KDSKESENKNFPNDSRKIIQTEFPNESKECDNIQF-LKDSKESENKNFPNDSRKIIQTEFPNESKECDNIQFLKDSKESNKYFLNESKKRKRDDHLNESKGSGNPNESRESETAEHLKEIGIDNPTKNDGIEIINRRSERLKTKPQISYNEEDNSLNKVVLNAHTIFNDVPNSFDEIQYRDDKSSWEEAINTELNAHKINNTWTITKRPENKNIVDSR WVFSVKYNELGNPIRYKARLVARGFTQKYQIDYEETFA and immunized in rabbits (CUSABIO Technology LLC).

### Immunocytochemistry

*Drosophila* wandering third instar larva body wall muscles were dissected in a manner previously described [4] calcium-free saline and fixed in 4% paraformaldehyde in 0.1M phosphate buffer, pH 7.2. Samples were washed and permeabilized in 0.1M phosphate buffer supplemented with 0.2% Triton X-100 (PBT) and incubated in a primary antibody overnight at 4°C. The samples were incubated with secondary antibodies at room temperature for two hours before mounting with Vectashield Mounting Media (Vector Laboratories Inc.). The primary antibodies used were: rabbit anti-aub, 1:1000 (see above), rat anti-aub, 1:2000 (see above), rabbit anti-Copia^Pol^ (see above) 1:1000 [1], rabbit anti-Copia^gag^, 1:5000 [1], rabbit anti-dArc1, 1:500 [4], rabbit anti-DLG, 1:40,000 [44], mouse anti-DLG, 1:200 (Developmental Studies Hybridoma Bank (DSHB), 4F3) and mouse anti-GFP, 1:500 (Developmental Studies Hybridoma Bank (DSHB), 4C9). Fluorescent conjugated anti-HRP and Alexa Fluor-secondary antibodies were used at a dilution of 1:200 (Jackson ImmunoResearch).

### Confocal Microscopy and Intensity Measurements

Confocal Z-stack images were acquired using a Zeiss LSM 800 microscope (40X Plan-Apochromat 1.30 NA DIC (UV) VIS-IR M27 and/or 63X Plan-Apochromat 1.40 NA DIC M27 oil immersion objectives). Images were quantified as previously described [45]. Briefly, volume and signal intensity from boutons of interest was measured using a protocol set up using Volocity software (Quorum Technologies Inc). HRP bound presynaptic volume was dilated and compared to DLG staining to obtain the postsynaptic volume. Protein levels were measured as fluorescence intensity and normalized to HRP volume of that bouton and data normalized to wild-type averages.

### RNA-FISH

RNA fluorescent in situ hybridization (FISH) was performed on *Drosophila* 3^rd^ instar larval body walls fixed in 4% PFA and incubated in 70% ethanol overnight at 4°C using custom Stellaris™ FISH Probes designed against *Aub* (NM_057386.5) by utilizing the Stellaris RNA FISH Probe Designer (Biosearch Technologies). *Aub* probes conjugated to CAL Fluor™ Red 590 dye (Biosearch Technologies, Inc.) were hybridized overnight at 37°C following the manufacturer’s instructions. The larval BWMs were co-stained with the neuronal marker HRP, washed and mounted with Vectashield Mounting Media (Vector Laboratories Inc.). The samples were imaged using an X-Light V2 spinning head confocal unit (Crest Optics) microscope equipped with the Orca-Fusion BT digital camera (Hamamatsu) and the Aura Light engine lasers (Lumencor) in widefield mode.

### RNA Isolation

The larval CNS and body wall muscles were isolated separately, and total RNA was isolated with the RNeasy Mini kit (QIAGEN). 10 µl of β-mercaptoethanol (Sigma-Aldrich) was added to 1 ml of buffer RLT (QIAGEN) and 350 µl of the buffer used to disrupt the tissue. The homogenate was cleared by centrifugation at max speed and transferred to a 2 ml safe lock sample tube. The samples together with the appropriate reagents were loaded onto the QIAcube connect (QIAGEN) and the pre-loaded RNA isolation protocol run. RNA integrity was determined by a Bioanalyzer (Agilent) while concentration was determined by a Qubit RNA assay (Invitrogen). The total RNA was shipped to Novogene Inc for library construction and sequencing.

### RNA-seq Analysis

Quality control of the reads was performed with FastQC. The reads were aligned to the Drosophila melanogaster reference genome using STAR [46]within a Snakemake [47] workflow. STAR was run with the following parameters: ––outFilterMultimapNmax 100 –– winAnchorMultimapNmax 200. TEtranscripts [48] was then used to quantify gene and transposable element (TE) abundances. TEtranscripts was run on the alignment files obtained from the STAR output using the following parameters: ––mode multi ––sortByPos ––format BAM. The GTF files for gene and TE annotations were supplied. Differential expression was determined using DESeq2 [49]. Volcano plots were created using the EnhancedVolcano package (https://bioconductor.org/packages/release/bioc/html/EnhancedVolcano.html). Both DESeq2 and EnhancedVolcano packages were run in RStudio.

### Alternative Splicing Analysis

Alternative splicing analysis was performed with the SpliceWiz package [50] in R (v4.4.2). In SpliceWiz, the default filter settings were used and edgeR was selected for the differential analysis. The coverage plots were obtained with the same SpliceWiz R package.

### Annotation Files

The Drosophila melanogaster reference sequences correspond to the toplevel sequences unmasked in the BDGP6.32.106 genome build from Ensembl. The fasta file with the reference sequences was downloaded from http://ftp.ensembl.org/pub/release-106/fasta/drosophila_melanogaster/dna/. The genomic feature annotation file in the General Transfer Format (GTF) for genes was downloaded from http://ftp.ensembl.org/pub/release-106/gtf/drosophila_melanogaster/. The GTF file for transposable element annotations was downloaded from https://labshare.cshl.edu/shares/mhammelllab/www-data/Tetranscripts/TE_GTF/.

### Total and small RNA Immunoprecipitation

Adult wild type *Canton-S* and *EGFP::aub^KI^* flies reared in RT were flash frozen five days after eclosion and stored in –80°C. The fly heads were isolated using a sieve placed on dry ice and the heads pulverized in a mortar placed on dry ice. The fly head powder was lysed using RIPA buffer (Abcam) supplemented with protease inhibitor cocktail (Roche), 1 mM PMSF, DNaseI (Qiagen), 2.5 mM MgCl_2_ and RNase inhibitor (Invitrogen) at 4°C. Lysates were rotated for 30 min and then centrifuged at 17000x g for 10 min at 4°C to remove tissue debris. The cleared lysates were mixed with equilibrated GFP-Trap agarose beads (Chromotek GmbH) and rotated end-over-end for 1 h at 4°C. Samples were then washed stringently with RIPA buffer. For small RNA isolation see below. Total RNA isolation was performed with the RNeasy mini kit (QIAGEN) in a QIAcube Connect instrument (QIAGEN) following pre-loaded protocols. For immunoblotting see below. beads were

### Small RIP sequencing (sRIP-seq) and analysis

Small RNA was eluted from the GFP beads-immune-complex with buffer RLT (Qiagen) supplemented with 2-mercaptoethanol and then purified using the miRNeasy mini kit (QIAGEN). Purified small RNA was used to generate precision small RNA libraries with the QIAseq miRNA UDI library kit (QIAGEN) following manufacturer’s recommendations. The prepared small RNA libraries were shipped to Novogene (Novogene Inc.) for single-end 50 (SE50) sequencing. The sRNAPipe in Galaxy was used for small RNA analysis [51]. The mapping was done allowing either 1 or two mismatches. The small RNAs were mapped to the Drosophila melanogaster BDGP6.46 version.

### Digital PCR (dPCR)

RNA samples were reverse transcribed into cDNA using the SuperScript IV First-Strand Synthesis System reaction (Invitrogen) following manufacturer protocol with RNase H digest. The dPCRs were multiplexed in 26K-partition 24-well or 8.5K-partition 96-well QIAcuity Nanoplates (QIAGEN) using a QIAcuity Digital PCR System (QIAGEN). For the reactions, either QIAcuity EvaGreen Master Mix or Probe Master Mix (QIAGEN) were used with the gene specific primer sets for *aub, Copia^full^, Copia^gag^*, Rpl32 or their probes (ThermoFisher or IDT). Data was processed in the QIAcuity Software Suite (QIAGEN), where absolute values (copies/µL) were obtained and normalized expression derived.

### Western Blotting

The GFP beads-immune complexes from RIP or IP were incubated directly with 4X protein loading buffer (Li-Cor) supplemented with 2-Mercaptoethanol (Sigma) and boiled at 95°C for 10 min. Proteins were separated in Mini-Protean TGX stain-free 4%–20% precast gels (Bio-Rad) under reducing and denaturing conditions. Proteins were transferred to an Immuno-Blot LF PVDF membrane (Bio-Rad) on a semi-dry Trans-Blot Turbo transfer system (Bio-Rad), blocked in Intercept Blocking Buffer (Li-Cor) and incubated with primary antibodies diluted in Intercept Antibody Diluent (Li-Cor) overnight at 4°C. Blots were washed, incubated with IRDye secondary antibodies (Li-Cor), washed again and finally imaged on a Li-Cor Odyssey CLx imaging system.

### Larval crawling assay

Vials of female and male flies of the appropriate genotypes were set up for mating. After 24 hours of laying eggs the adults were removed and the vials transferred from 25℃ to 29℃ incubators to allow for optimal transgene expression driven by Gal4, the yeast transcription factor. Wandering third instar larvae were collected and washed with deionized H_2_O. Larval movement was recorded on agar molded in weigh dishes in a manner previously described [52] and analyzed using a modified a freeware version of the ImageJ Plugin wrMTrck (ImageJ: http://imagej.nih.gov/ij/) and wrMTrck: http://www.phage.dk/plugins/wrmtrck.html), to extract relative speed and distance travelled. Data was analyzed and graphed with GraphPad Prism version 10.5.0 for Windows (GraphPad Software).

### Adult Locomotion Assay

Analysis for climbing ability was determined using a standard protocol [53]. This assay scores the flies’ ability to climb over their lifetime and analyzes 50 males from every genotype.

Climbing indices obtained were analyzed using GraphPad Prism version 10.5.0 for Windows (GraphPad Software) and climbing curves were fitted using non-linear regression. Comparisons were performed at 95% confidence interval with a P-value threshold of less than 0.05 considered significant.

### Adult *Drosophila* Aging Assay

Survival analysis was performed following a previously described protocol [53]. Briefly, crosses of each genotype were made from virgin females and males and a time matched cohort of at least 200 flies were collected. The flies were pushed to new food every other day and were considered dead when they did not display movement upon agitation. Survival curves were graphed, and the curves were compared using the log-rank (Mantel-Cox) test using GraphPad Prism version 10.5.0 for Windows (GraphPad Software).

### Quantification and Statistical Analysis

Statistical analysis for single comparisons was performed using a Student’s t test while comparisons across multiple experimental groups utilized a one-way analysis of variance (ANOVA) with the appropriate post hoc test. ∗, p < 0.05; ∗∗, p < 0.001; ∗∗∗, p < 0.0001. Raw data files were processed with Microsoft Office Excel software (Microsoft Inc.) and data analysis for statistical significance utilized GraphPad Prism version 10.5.0 for Windows (GraphPad Software).

## Declarations

### Availability of data and materials

The datasets supporting the conclusions of this article are included within the article and its additional files.

### Competing interests

The authors declare that there are no financial and non-financial competing interests that would result in conflict of interest.

### Funding

This study was funded by a National Institutes of Health grant R01NS112492 grant to TT.

### Author’s contributions

Conceptualization: TT and PGM. Methodology: PMG, DO, GA, JG, AM and TT. Investigation: PGM, DO, GA, JG, MZ, AO, AT, AM and TT. Visualization: PGM, DO, and TT. Formal analysis: PGM, DO, GA and TT. Supervision: TT. Resources: TT and PGM. Writing—original draft: TT, PGM and AM. Writing—review and editing: PGM, GA, JG, MZ, AM and TT.

## List of abbreviations

BWM: body wall muscle
CNS: central nervous system
GFP: green fluorescent protein
KD: knockdown
LNMJ: larval neuromuscular junction
RFP: red fluorescent protein
RIP: RNA immunoprecipitation
TE: transposable element

## Acknowledgements

We are thankful to Dr. Tatjana Trcek for generously sharing the EGFP::aub^KI^ fly line, and to the Bloomington and Vienna Drosophila Stock Centers for the fly lines. FlyBase (NIH P40OD018537) for managing the fly database.

## Figure Legends

**Figure S1.**
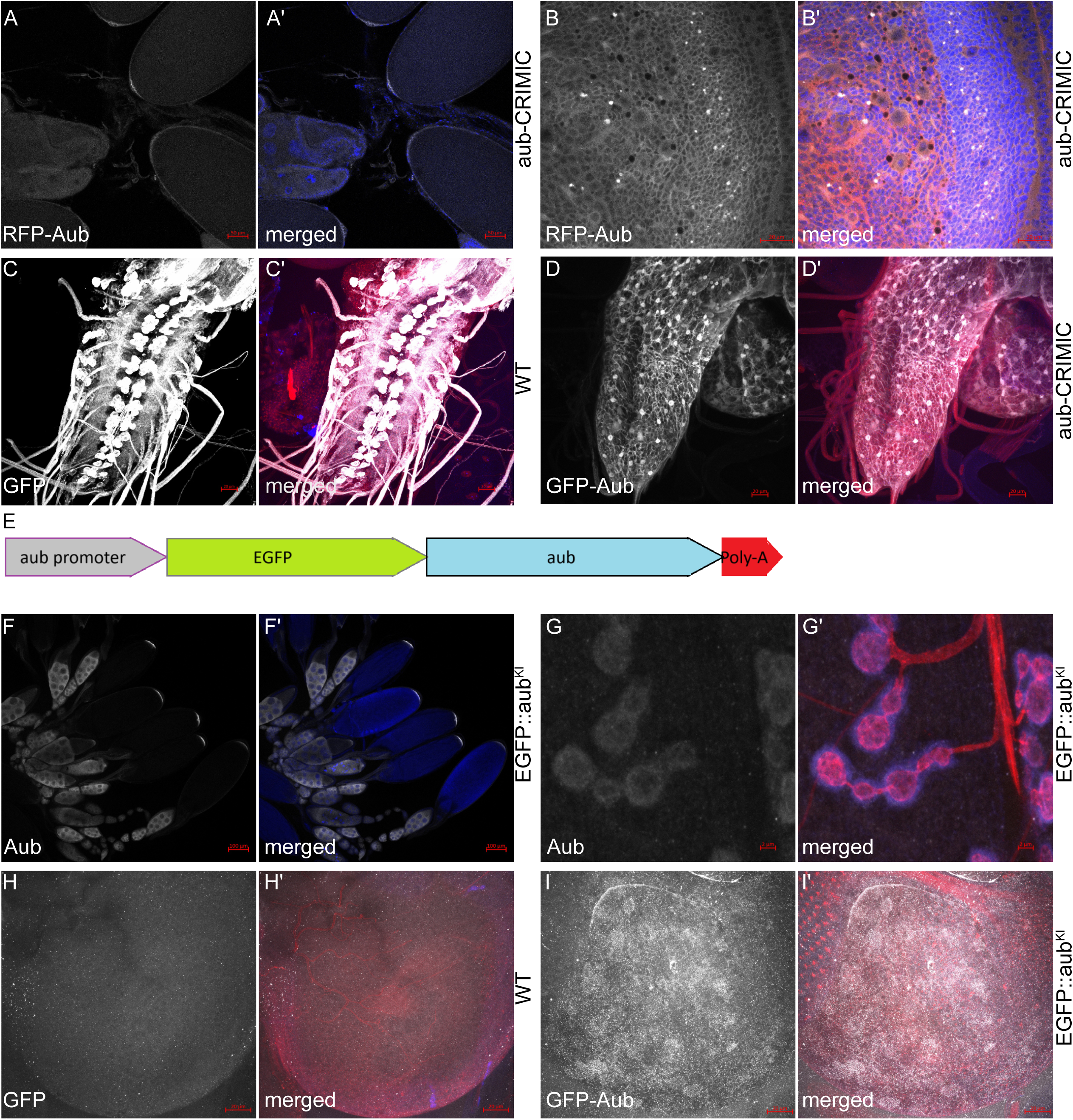
Aub is enriched in germline and somatic tissues. (A) Dissected adult ovary tissue of the *aub*-CRIMIC fly line, with RFP indicating the endogenous *aub* expression pattern in ovaries; (A’) merged image of the orthogonal projection with DAPI. Scale bars, 50 μm. (B) Dissected third instar larval brain lobes of the *aub*-CRIMIC fly line, with RFP indicating the endogenous *aub* expression pattern in a select neuronal population; (B’) merged image of the orthogonal projection with HRP, DAPI and RFP. Scale bars, 20 μm. (C) Dissected third instar larval VNC of a C380-Gal4/GFP fly line (note as “WT” in panel), expressing GFP in motor neurons via the motor neuron specific C380-Gal4 driver; (C’) merged image of the orthogonal projection with HRP, DAPI and RFP. Scale bars, 20 μm. (D) (D) Dissected third instar larval VNC of the *aub*-CRIMIC fly line, with GFP indicating the endogenous *aub* expression pattern in a select motor neurons; (D’) merged image of the orthogonal projection with HRP, DAPI and RFP. Scale bars, 20 μm. (E) Diagram of the *EGFP::aub^KI^* fly line locus showing the knocked-in EGFP under the control of the endogenous aub promoter. Scale bars, 20 μm. (F) Dissected adult ovary tissue of the *EGFP::aub^KI^* fly line, with GFP indicating the endogenous *aub* expression pattern in ovaries; (F’) merged image of the orthogonal projection with DAPI. Scale bars, 100 μm. (G) Dissected third instar LNMJ of the *EGFP::aub^KI^* fly line, with GFP indicating the endogenous *aub* expression pattern at the LNMJ; (G’) merged image of the orthogonal projection with DAPI. Scale bars, 2 μm. (H) Dissected third instar larval brain lobes of the WT (Canton-S) control fly line, with anti-GFP staining indicating lack of background GFP signal; (H’) merged image of the orthogonal projection with neuronal tissue staining (HRP). Scale bars, 20 μm. (I) Dissected third instar larval brain lobes of the *EGFP::aub^KI^* fly line, with GFP-positive staining indicating the endogenous *aub* expression pattern; (I’) merged image of the orthogonal projection with neuronal tissue staining (HRP). Scale bars, 20 μm.

**Figure S2.**
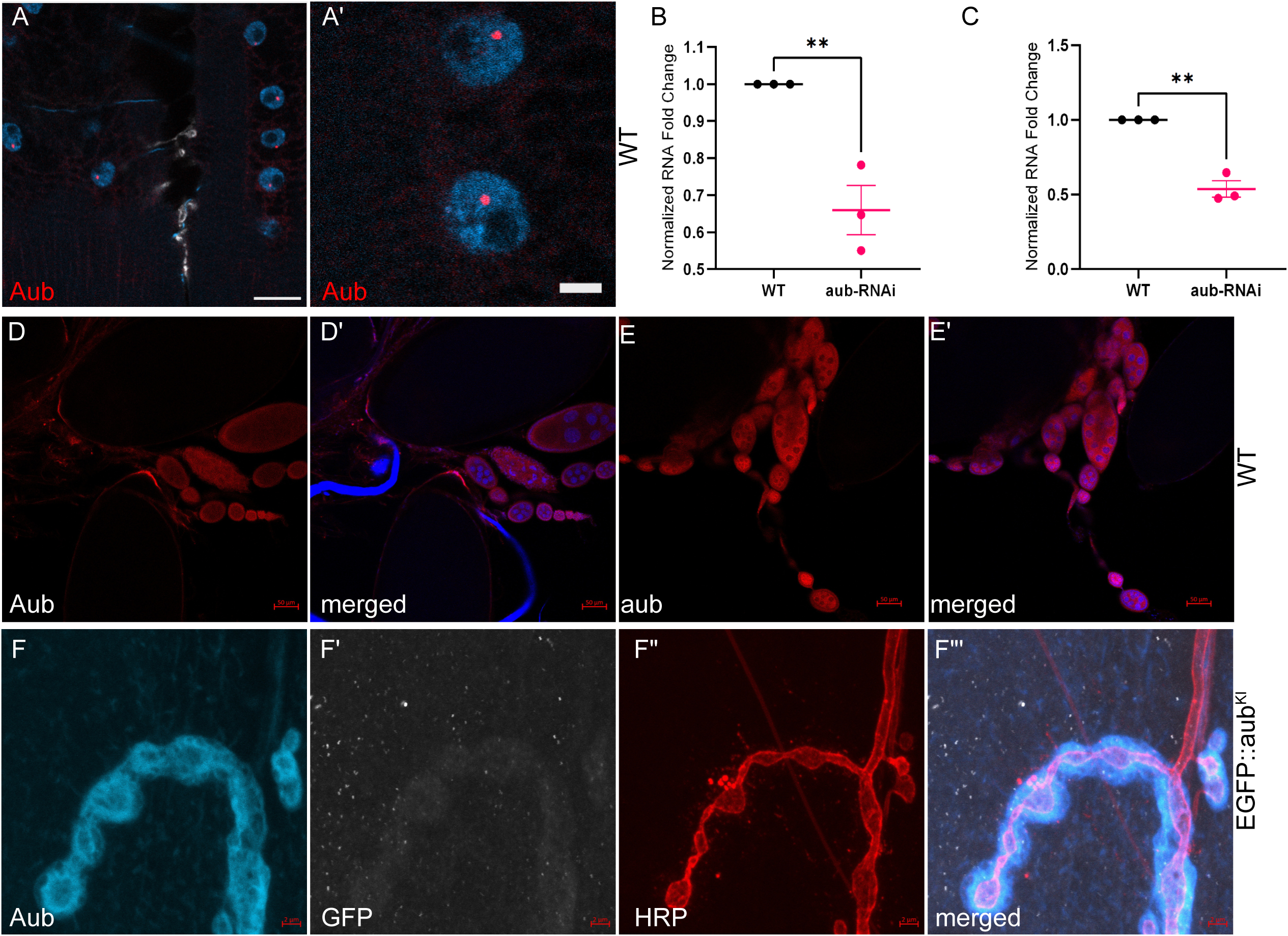
Aub protein is enriched in neurons and tissue around the larval NMJ. (A) Dissected third instar larval BWM of a WT (Canton-S) control fly line, stained for *aub* mRNA using RNA fluorescence in situ hybridization (FISH); magenta, *aub* mRNA probe staining; blue, DAPI nuclei staining. Scale bar, 20 μm. (A’) Magnified nuclei as in (A). Scale bar, 5 μm. (B) Quantification of *aub* expression in larval CNS tissue via dPCR; comparison of WT (C380-Gal4/Canton-S) control and motor neuron specific (C380-Gal4 driven) presynaptic *aub* knockdown fly lines. N = 3. (C) Quantification of *aub* expression in larval BWM tissue via dPCR; comparison of WT (C57-Gal4/Canton-S) control and muscle specific (C57-Gal4 driven) postsynaptic *aub* knockdown fly lines. N = 3. (D) Dissected adult ovary tissue from the WT (Canton-S) control fly line, stained for *aub* protein expression using antibody generated in rat against an AUB peptide antigen (see Methods). Scale bar, 50 μm. (E) Dissected adult ovary tissue from the WT (Canton-S) control fly line, stained for aub protein expression using antibody generated in rabbit against an AUB peptide antigen (see Methods). Scale bar, 50 μm. (F) Dissected third instar LNMJ of the *EGFP::aub^KI^* fly line, stained with anti-AUB antibody; (F’) anti-GFP staining indicating EGFP-AUB; (F”) HRP staining of the motor neurons of the LNMJ; (F’”) merged image of the orthogonal projection, showing AUB immunoreactivity and motor neuron staining (HRP). Scale bars, 2 μm. “ns” p≥0.05, * p<0.05, ** p<0.01, *** p<0.001, and **** p<0.0001. Error bars are SEM.

## Notes

### Competing Interest Statement

The authors have declared no competing interest.

